# Context dependency of nucleotide probabilities and variants in human DNA

**DOI:** 10.1101/2021.07.22.453351

**Authors:** Yuhu Liang, Christian Grønbæk, Piero Fariselli, Anders Krogh

## Abstract

**Background:** Genomic DNA has been shaped by mutational processes through evolution. The cellular machinery for error correction and repair has left its marks in the nucleotide composition along with structural and functional constraints. Therefore, the probability of observing a base in a certain position in the human genome is highly context-dependent.

**Results:** Here we develop context-dependent nucleotide models. We first investigate models of nucleotides conditioned on sequence context. We develop a bidirectional Markov model that use an average of the probability from a Markov model applied to both strands of the sequence and thus depends on up to 14 bases to each side of the nucleotide. We show how the genome predictability varies across different types of genomic regions. Surprisingly, this model can predict a base from its context with an average of more than 50% accuracy. For somatic variants we show a tendency towards higher probability for the variant base than for the reference base. Inspired by DNA substitution models, we develop a model of mutability that estimates a mutation matrix (called the alpha matrix) on top of the nucleotide distribution. The alpha matrix can be estimated from a much smaller context than the nucleotide model, but the final model will still depend on the full context of the nucleotide model. With the bidirectional Markov model of order 14 and an alpha matrix dependent on just one base to each side, we obtain a model that compares well with a model of mutability that estimates mutation probabilities directly conditioned on three nucleotides to each side. For somatic variants in particular, our model fits better than the simpler model. Interestingly, the model is not very sensitive to the size of the context for the alpha matrix.

**Conclusions:** Our study found strong context dependencies of nucleotides in the human genome. The best model uses a context of 14 nucleotides to each side. Based on these models, a substitution model was constructed that separates into the context model and a matrix dependent on a small context. The model fit somatic variants particularly well.

## Background

The evolution of species can be followed in chromosomal DNA, which has undergone mutations and selection, and mutational processes have been essential for the development of life on earth. On the other hand mutations need to be controlled, because if an essential gene is mutated it may result in severe disease or loss of viability. This balance between plasticity and stability is important for sustaining stable life forms [1]. The question we ask in this study is, how this balance is reflected in the local sequence properties of human DNA and how the sequence context affects mutations. More precisely, we consider models of mutability that depend on the sequence context of e.g. *k* bases on each side of the position in question.

It is well known that the sequence context influences mutational processes. For instance, the mutation of C to T is much more common in CpG dinucleotides than in other contexts in the human genome[2, 3], and previous studies have reported that the immediate neighbouring bases (up to a 7 base context) influence mutation rates [4, 5, 6, 7]. Another study showed point mutations can be affected by sequence motifs [8]. The cellular machinery includes components for maintaining genome integrity, such as DNA repair mechanisms, which result in mutational biases[9, 10] and other processes may lead to other biases. These mechanisms together govern the intrinsic mutability. Following [11], we use the term mutability rather than mutation rate, because we are not considering the detailed evolutionary process and there is no time in our models, although the same ideas are easily applicable to estimation of context sensitive mutation rates.

Models of mutability can be estimated from observed variants by simply estimating the probability of a mutation given a context. However, such models are estimated from fairly small and biased sets of variants without utilizing the mutability foot-print in the genome. Here we propose to split the context dependent mutability into a nucleotide distribution and a variant part. The nucleotide distribution can be estimated from the whole genome and the variant part from variants, thereby allowing the two parts to have different context sizes. Due to the size of the human genome, the context dependent nucleotide distribution can be estimated from a much larger context than the variant part. The variant part can depend on a *smaller* context and can thus be estimated from a small number of variants.

In the first part of the paper, we focus on estimation of the probability of observing a base in the genome, given a context. One measure to quantify the context sensitivity is predictability. In a random sequence of nucleotides with no context sensitivity, we would only be able to predict a given base with an accuracy of 25% (random guessing), so this is the lower boundary of predictability. However, due to the mutational biasses discussed above and the repetitive nature of genomes, we would expect that a genome is more predictable than a random sequence. We show that a human genomic base can be predicted with an average of 51% using our most sophisticated model.

In the second part of the paper, we estimate a mutability model based on the context dependent nucleotide distribution found. For a fixed context dependent nucleotide distribution model, we show that the mutability is not very sensitive to the context size of the variant part. We compare to a simple mutability model conditioned on a 7 base context as in [5] and show that they differ between different types of mutations.

Knowledge of the background probability is important for a lot of models and the models described in this work can form a basis for other modelling efforts in the future. It has been shown, for instance, that a high-order Markov model can improve motif discovery over a simple background model [12]. Similarly our models of mutability can be useful in future studies of mutations in disease, where the mutability can be used to e.g. identify unexpected mutations.

## Results

### Context modeling of the human genome

In our first model, the Central model, (Figure 1), we simply estimate the conditional probability of a nucleotide given *k* bases to each side. For base *x*_*i*_ at a genomic position *i* these probabilities are written as

**Figure 1.**
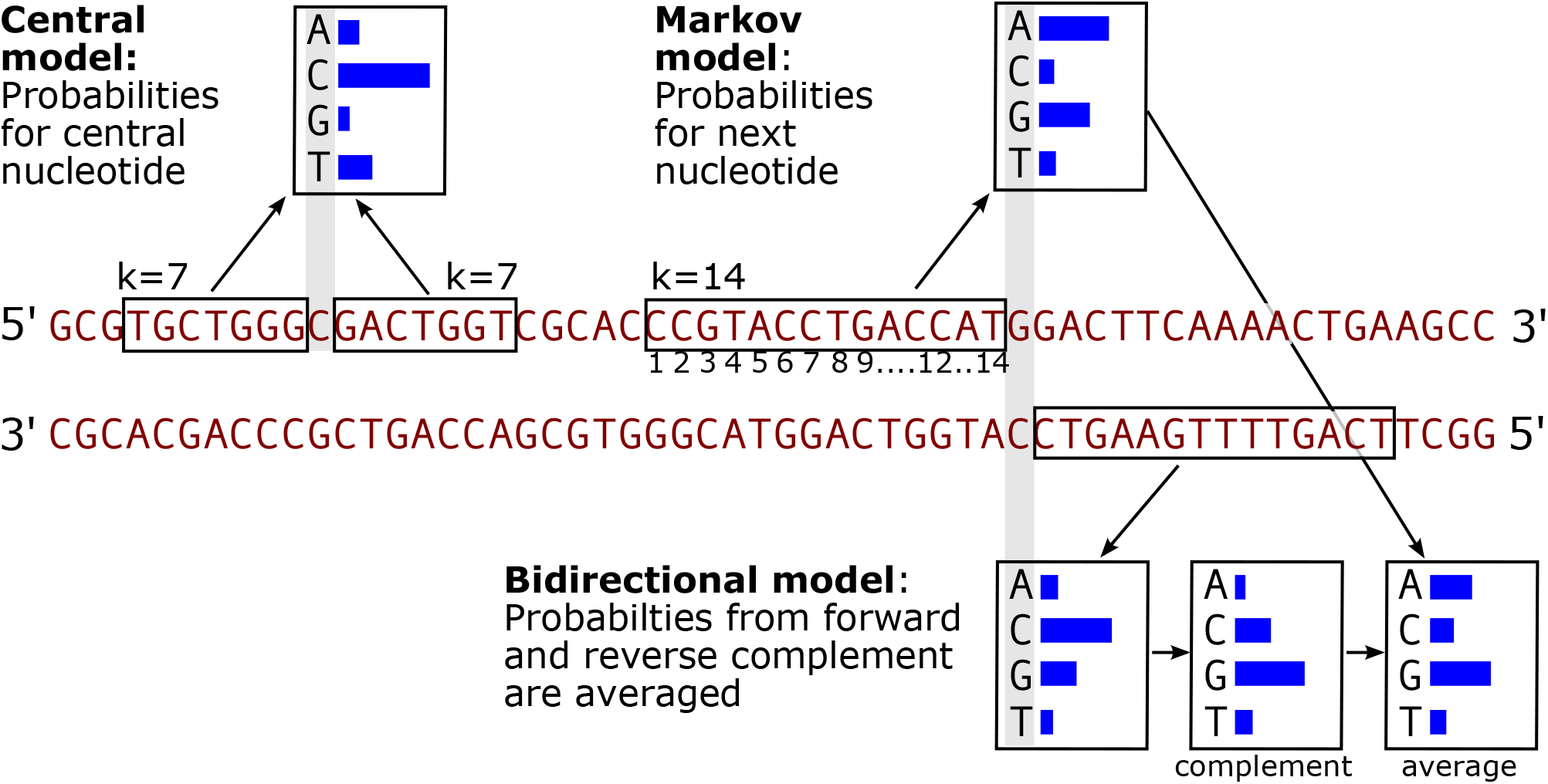
Illustration of the three models used. A DNA sequence is shown with the complement sequence below. The blue histograms illustrate nucleotide probabilities. The central model with *k* = 7 (upper left) predicts the base in the middle from the adjacent nucleotides in the boxes to the left and right. For this illustration, C has highest probability, which happens to coincide with the correct nucleotide at the position. The Markov model (top right) of order *k* = 14 predicts a nucleotide from the previous 14. In this example A has highest probability although G is the actual reference probability. The bidirectional model (bottom right) use the same model on the reverse complement strand. In this example C has the highest probability, which coincides with the complement base at the position. The probabilities are translated to the direct strand and averaged with the forward model.

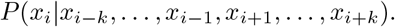

They are estimated from the genomic frequencies of the 4 possible (2*k* + 1)-mers of the given context. A *k* = 3 model corresponds to a neighbourhood of 7 as used in [5], and we use this model as our baseline. Since we are estimating frequencies from all positions on both strands, they are automatically strand symmetric.

One can use other values of *k* as long as a model can be reliably estimated. As the 4 probabilities sum to one, there are 3 * 4^2*k*^ free parameters in the model, so the *k* = 3 model has around 12,000 free parameters, which can easily be estimated from the 6 billion sites of the two strands of the human genome. A *k* = 7 model has approximately 0.8 billion free parameters, and is thus the upper limit of what we can hope to reliably estimate for a genome like the human. Even with *k* = 7 there are many contexts that occur only once or very rarely. To avoid over-fitting, we have used an interpolated Central model in which a model of order *k* is used to regularize a model of order *k* + 1 and so on (see Methods). For our second model, we have used a central model with *k* = 7 and interpolated from *k* = 4.

A Markov model of order *k* yields probabilities of the four bases conditional on the *k* previous bases. A Markov model also can be used to estimate from both strands, as above, which means that for base *i*, it can give two different probabilities: *P* (*x*_*i*_|*x*_*i*−1_, …, *x*_*i*−*k*_) on the direct strand and 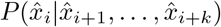 on the opposite strand, where 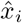 means the complementary base to base *x*_*i*_. Note that these models are estimated from both strands as the central models, which means that a model estimated using a 5’ context is identical to the complementary of a model estimated using a 3’ context and therefore, without loss of generality, we always assume 5’ models.

Our third model is a bidrectional Markov model (Figure 1) of order *k* = 14, interpolated from *k* = 8. It is called bidirectional, because we use the *average* between the probability of *x*_*i*_ from one strand and the probability of 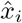 from the opposite strand as explained above. Note that this model with *k* = 14 has the same number of free parameters (3 * 10^14^) as the central model with *k* = 7 described above, because both use 14 bases as context. However, the bidirectional Markov model actually uses a context of 28 bases for prediction, because of the averaging over the two directions. This model is called BM14 in the following.

We have developed a program written in C that implements these different models. Instead of saving counts for each context, it dynamically calculates the count based on a Burrows-Wheeler encoded genome [13] to save memory. The performance of our models can be evaluated by the accuracy, which is the fraction of positions, where the most probable base given the context equals the actual base in the reference genome. The accuracy on the human genome is shown in Figure 2 for the different models mentioned above (Supplementary Table S1, S2).

**Figure 2.**
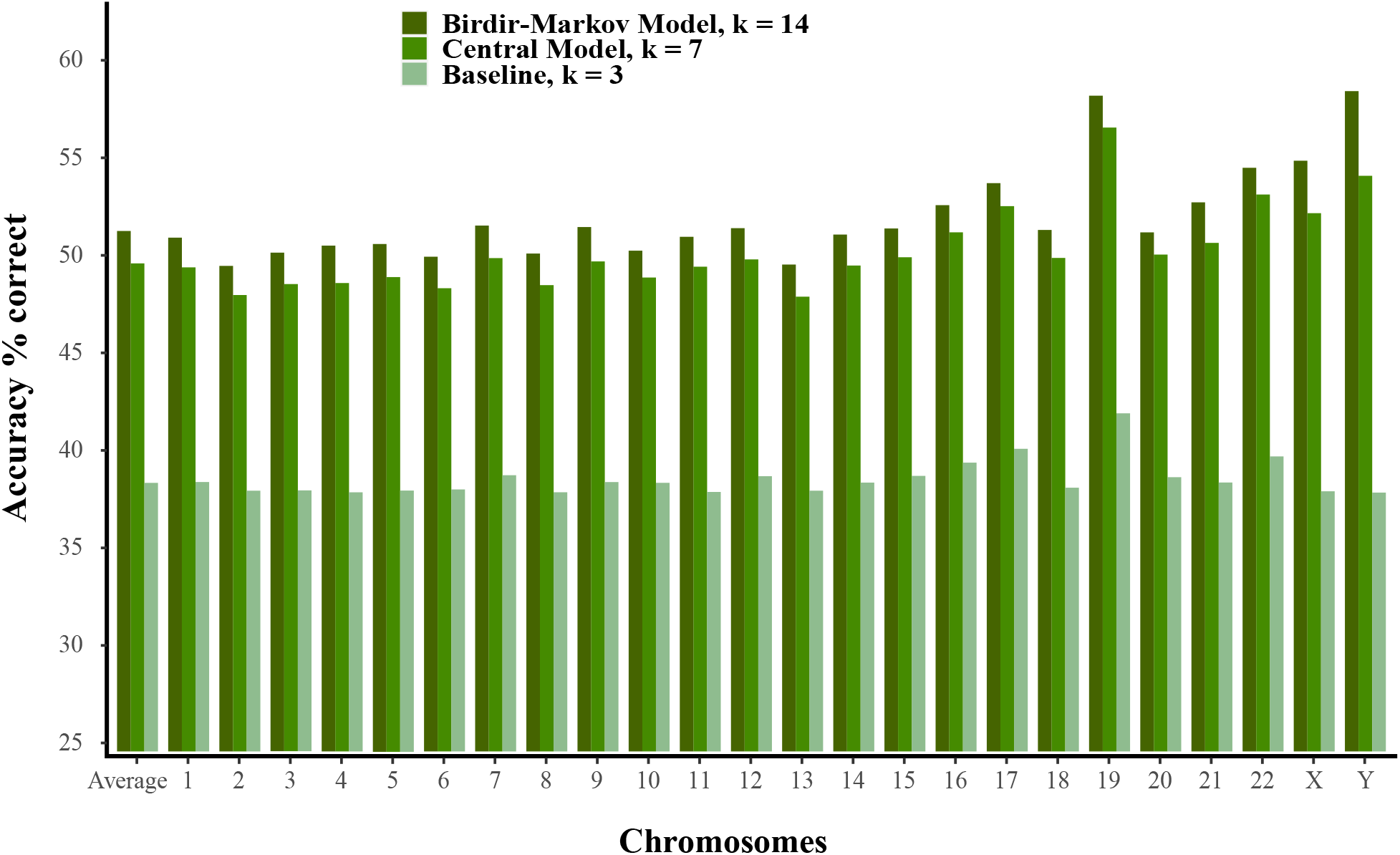
Prediction accuracy for the three models. Baseline with *k* = 3, Central model with *k* = 7 and the bidirectional Markov model with *k* = 14 (BM14). The bar-plot shows accuracy for each chromosome and average accuracy on the whole genome. Results using nucleotide-based cross-validation.

For the baseline model there is a strong correlation between the GC content and the accuracy on each chromosome. In Supplementary Table S3, we show GC content [14] with the accuracy and find a Pearson correlation of 0.90 for the baseline model with the lowest accuracy of around 38% for Chromosome 2–6 that has GC content of 38–40% and the highest accuracy of around 42% for chromosome 19, which has the highest GC content of 48%. For the *k* = 7 central model and BM14, the picture is less clear. Although they have correlations of 0.70 and 0.53 with GC content, the two chromosomes with the best prediction accuracy are chromosome 19 (GC 48%) and chromosome Y (GC 40%) at opposite ends of the GC scale.

For estimating the performance shown in Figure 2, we have used leave-one-out cross-validation at the nucleotide level. It means that when estimating the probabilities for a given site in the genome, that site is excluded in the counts for model estimation. Because the k-mers overlap, one may argue that it is not proper cross-validation, but more fulfilling a minimum requirement that the site itself should not be used for estimating the model. Therefore we have also done a chromosome-based cross-validation for comparison and calculated the overall accuracies for each chromosome using a model estimated from the *other* chromosomes. The difference between nucleotide-based and chromosome-based cross validation is only 0.5 percentage points (p.p.) on average, but for the Y chromosome, it is more than 3 p.p. (Supplementary Table S1, S2 and Supplementary Figure S1). Chromosome Y is known to differ from other chromosomes by being more heterochromatic and contain mostly repetitive regions [15], and therefore the model performs poorly on this chromosome when estimated only from other chromosomes.

With interpolation it is in principle possible to go beyond *k* = 14, because for contexts with zero counts, the probabilities are equal to a lower order estimate, so it should adapt without over-fitting. We have not explored higher *k* so much, but in Supplementary Figure S2, we have run the bi-directional Markov model from *k* = 10 to *k* = 20 for different values of the interpolation constant described in Methods. The figure shows results for chromosome 20 and the model estimated from all the other chromosomes. Up to *k* ≃ 14 the models steeply improve and are almost insensitive to the interpolation constant. Above *k* = 14 we still see a monotonous improvement that seems to level off at around 52% for the best model. Chromosome 20 was chosen for this experiment, because it is small and has a prediction accuracy similar to the average for the BM14 model. It clearly shows that interpolation improves the model although not by a great deal for *k* < 14. Importantly, interpolation at any strength ensures that zero counts do not occur, which would otherwise result in undefined probabilities.

The predictive performance of BM14 on different regions in the human genome is shown in Figure 3. As expected, the model predicts repetitive sequences very well with an overall accuracy of 64%, but there are quite large differences between different types of repeats. The most common type of repeat in the human genome, the ALU sequences, is 87% correctly predicted, whereas LINE1 for instance is only at 63% (Supplementary Table S4). These differences are most likely due to differences in conservation of the different types of repeats.

**Figure 3.**
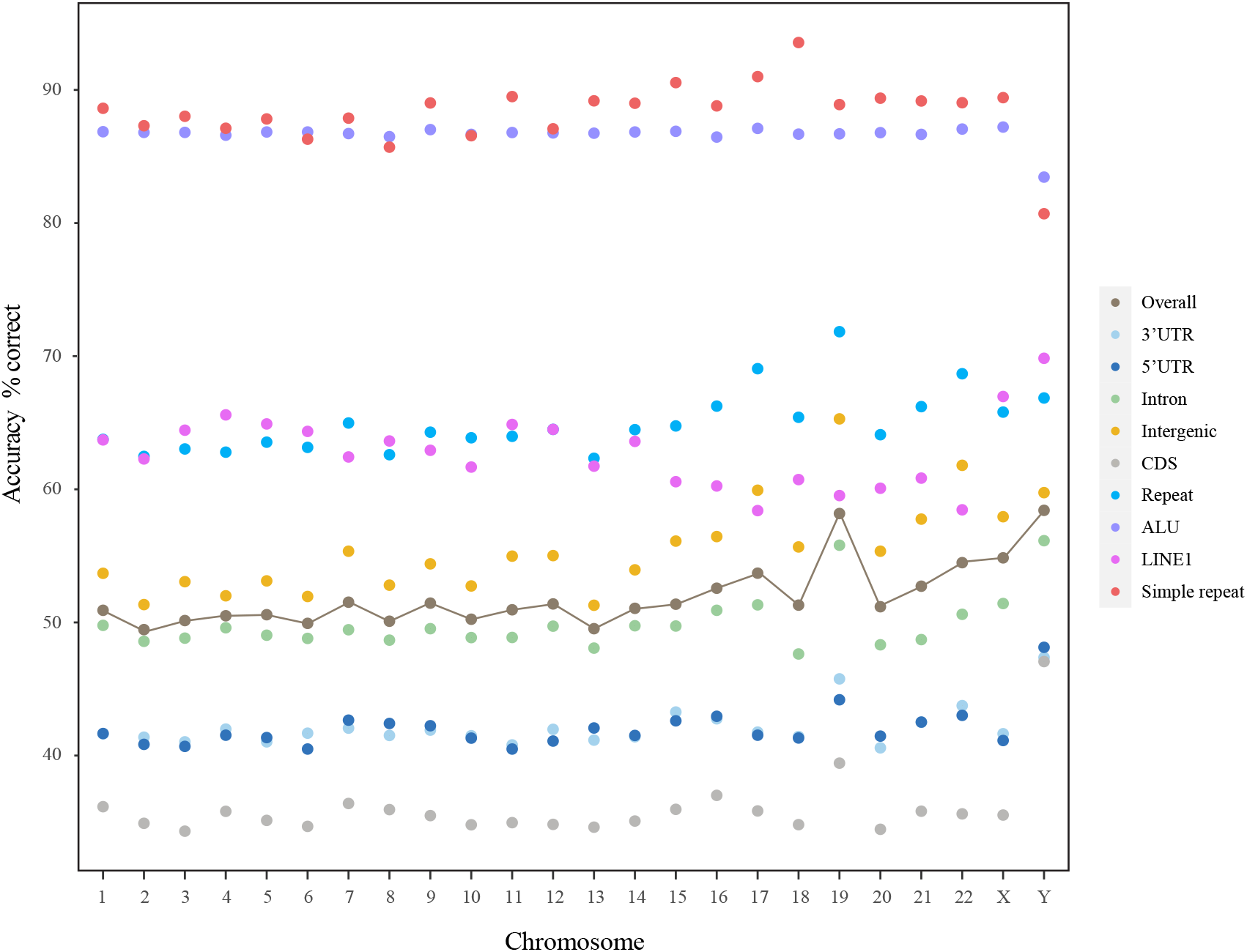
Prediction accuracies for BM14 in different regions across all chromosomes. The accuracy for different features on the chromosome 1 to Y, is indicated by colored dots. The line shows the overall accuracy for each chromosome.

The probability of the nucleotide in the reference genome given its context varies throughout the genome. The density of this probability, which we call the reference probability, is shown for different genomic regions in Figure 4. For each feature except for CDS there are two peaks of which one is due to repeats. However, in positions where the reference probability is above 0.4, repeats account for a large proportion compared to other features. (Supplementary Table S5).

**Figure 4.**
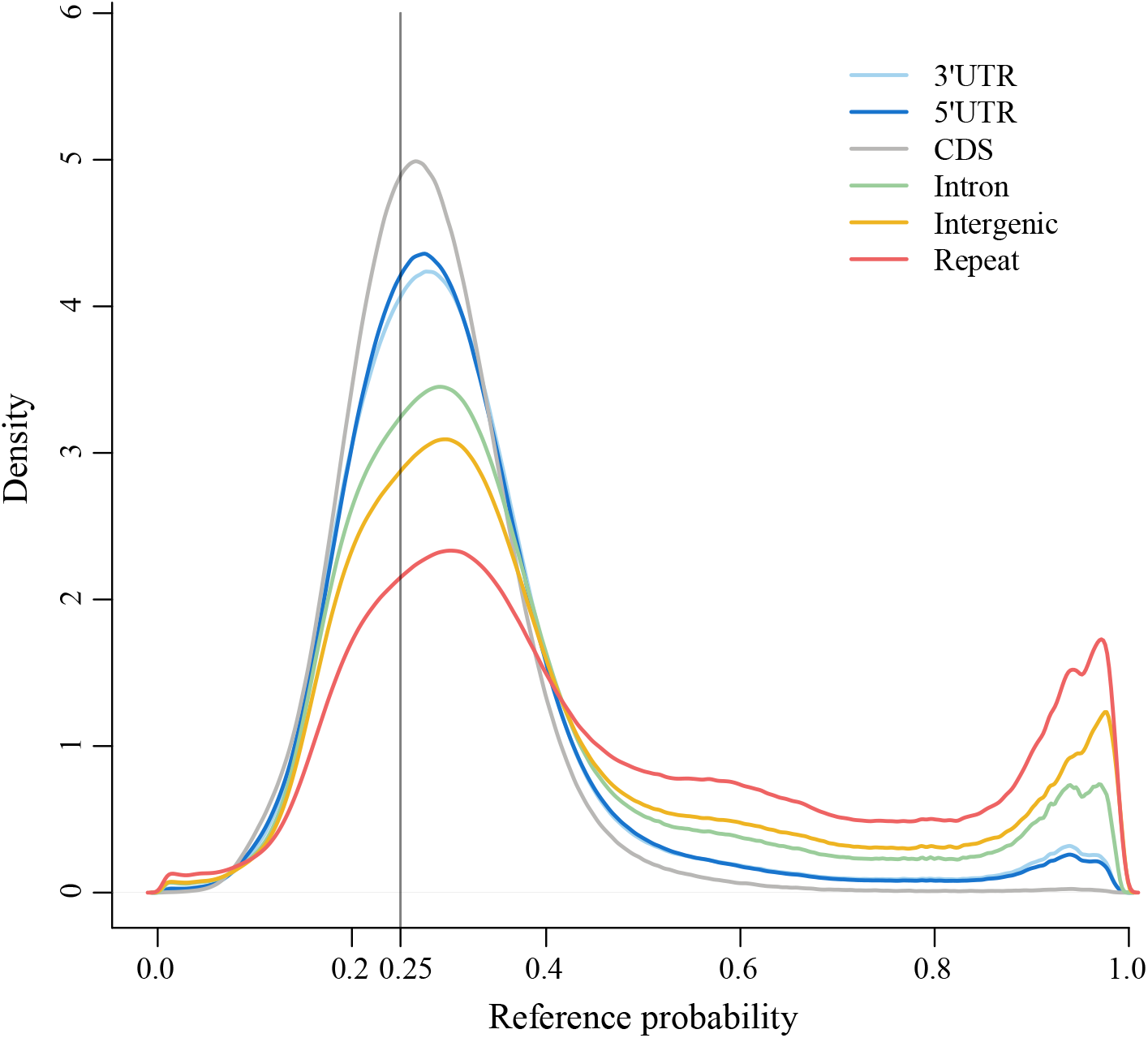
Density profile of reference probabilities in different genomic regions obtained with BM14.

To further elucidate the predictability across different regions, we show in Figure 5 the reference probabilities across human 3’ and 5’ splice sites that averaged over all introns annotated in Chromosome 1 (Chr1). The probability shows a large jump from a level of almost random prediction (∼ 0.28) in the coding region to a fairly high value (∼ 0.36) in the intron. The conservation plot in the same figure presents an opposite trend.

**Figure 5.**
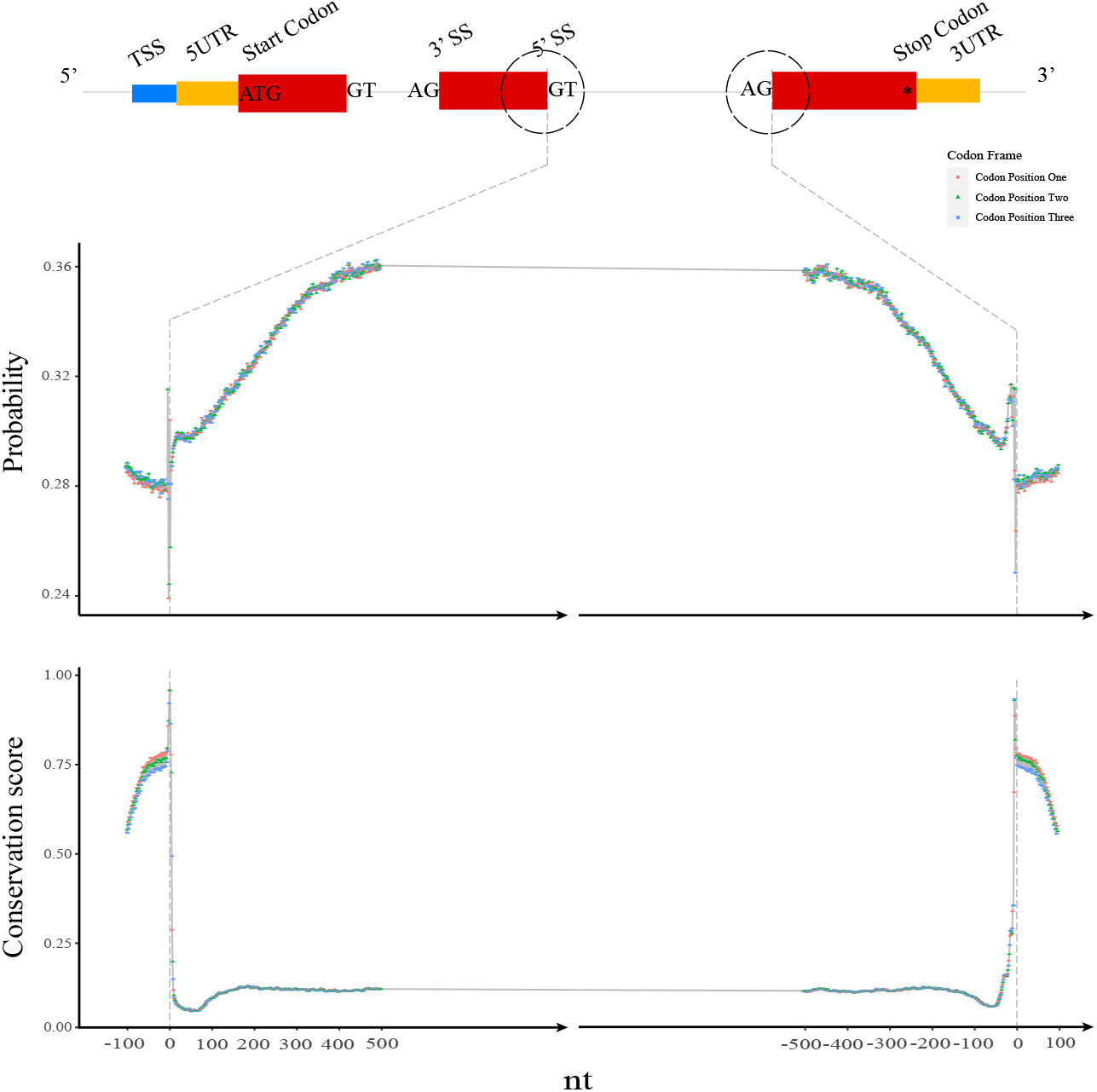
Probabilities (top) and conservation score (bottom) of reference bases across 3’ and 5’ splice sites. The probabilities of the reference bases by BM14 were averaged for each position for the first/last 100 nt in coding sequence and 500 nt in introns. The conservation score is PhastCons100Way from the UCSC browser.

To test whether the model can be improved for non-repeat regions, we estimated a restricted model from everything *outside* coding regions and repeats. There is little difference between the restricted model and the full one in terms of prediction accuracy or reference probability as seen in (Supplementary Figure S3) and we did not analyze this model further.

We briefly examined the performance of a bidirectional Markov model on some other species. Because of the smaller genome sizes, we used an interpolated bidirectional Markov model of order *k* = 10 in this analysis. The density plot of the reference probabilities (Supplementary Figure S4A) shows that a single main peak occurs for human and *E*.*coli* genomes. *A. thaliana, C. elegans* and *S. cerevisiae* have two peaks. The peak towards low probability is enriched in coding sequence as can be seen from Supplementary Figure S4B, where the density is plotted separately for CDS regions and other regions. In positions where the reference probability is above ∼ 0.55, the density of human is higher than that of other species, which is most likely caused by repeats in human genome.

In the other eukaryotic genomes the prediction accuracy of the models were 45% for *C. elegans*, 40% for *A. thaliana*, and 38% for *S. cerevisiae*.

### Variants

We next evaluated BM14 on variant datasets. We assume that our models are valid for all genomes, and variants found in population studies, such as the 1000 Genomes Project (1KGP) [16], should be predicted with the same accuracy as the corresponding positions in the reference genome. We identified ∼ 73 million biallelic single nucleotide polymorphisms (SNPs) in the 1KGP. The probability of the reference (Pref) was plotted against the probability of the alternative (Palt) shown in Figure 6 for the *k* = 7 central model and BM14. The latter shows a larger concentration of sites in the middle of the plot. Note the unexpected asymmetry between the corners at Pref≃1 and Palt≃ 1 for both models.

**Figure 6.**
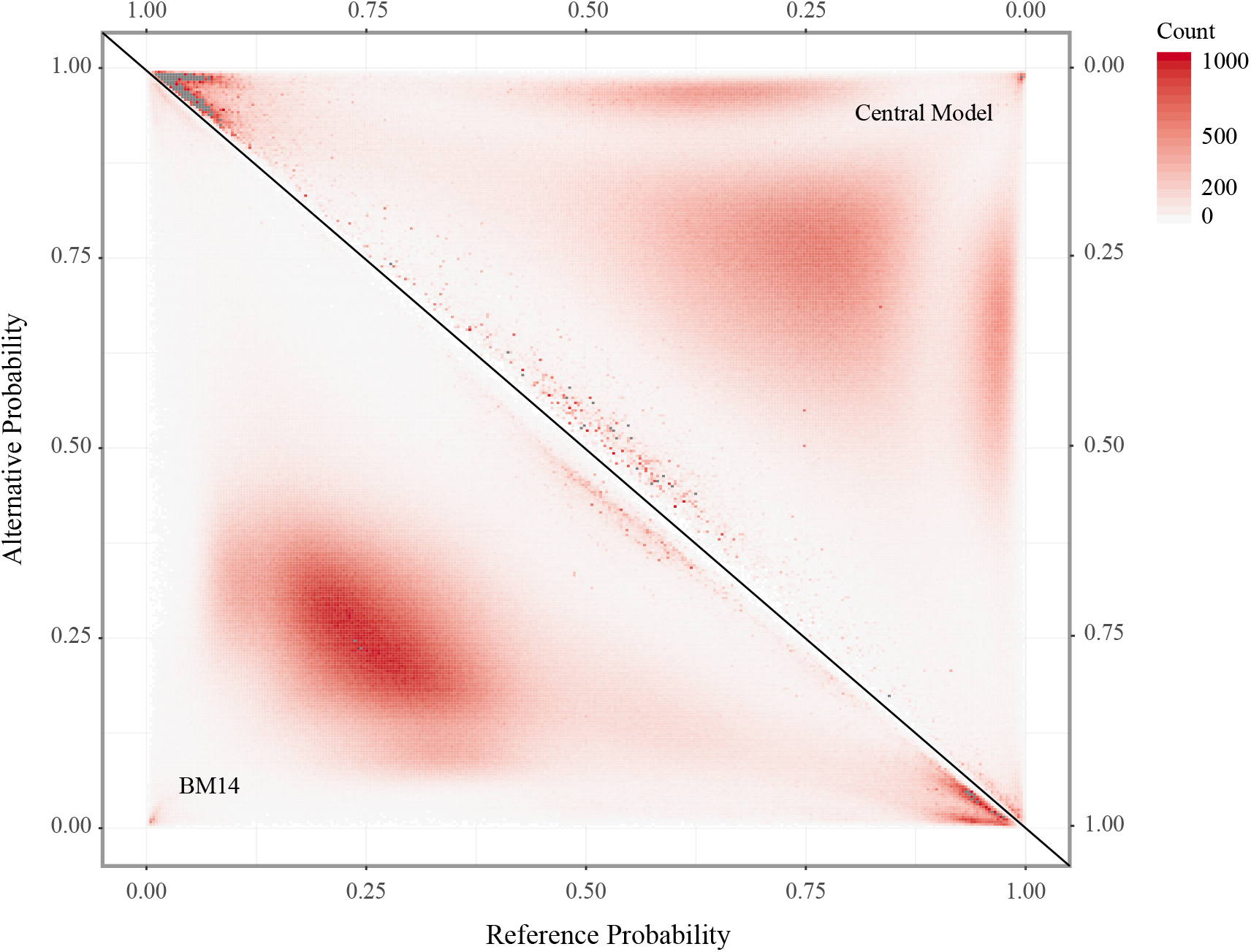
Triangle plot for probabilities of Ref-Alt alleles. Probabilities of reference and alternative alleles were estimated by the *k* = 7 central model (upper right triangle) and the *k* = 14 bidirectional Markov model (BM14, lower left triangle) on SNPs from the 1000 Genomes Project.

This asymmetry is also reflected in the fact that the reference allele had the highest probability in 38.82% of cases and the alternative allele in only 24.20% for BM14. The density plot of Pref-Palt in Figure 7A also shows a peak near 1 when all SNPs are used. However, when rare SNPs are ignored, the right peak decreases in size and a peak in the left side of the plot appears and the density becomes symmetric when only including SNPs with allele frequency above 20%. The far majority of SNPs with a reference probability higher than 0.875 in the 1KGP dataset belong to repeats.

**Figure 7.**
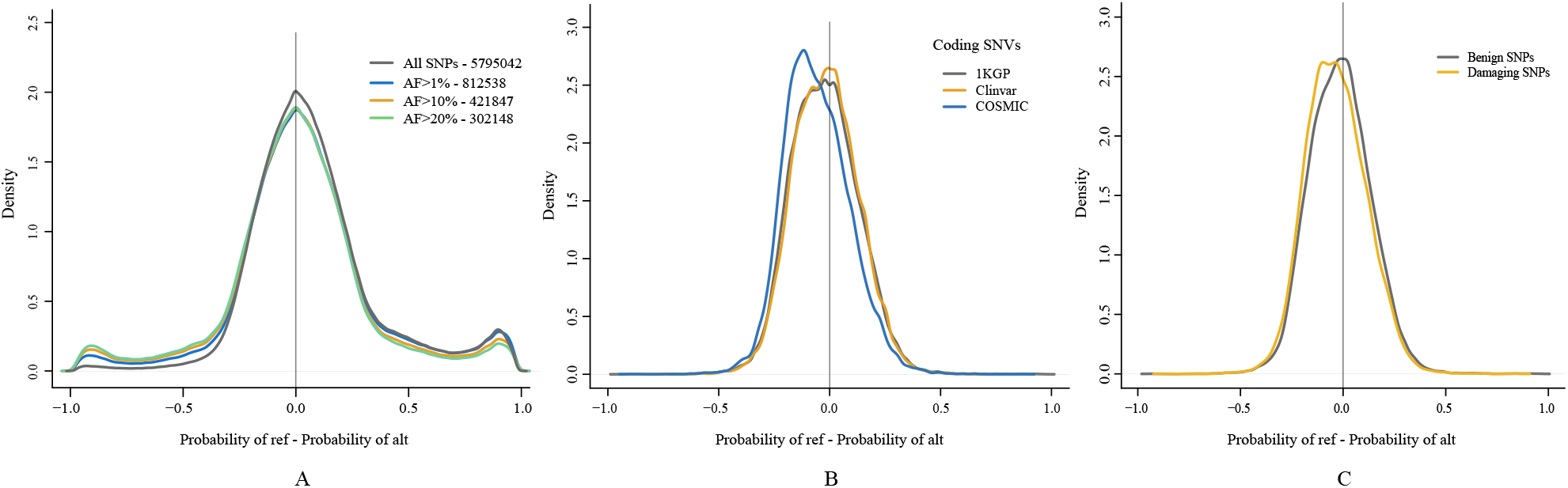
Density profiles of Pref - Palt for SNPs on Chromosome 1. A. SNPs from 1KGP. The different lines represent SNPs with allele frequencies greater than 0, 0.01, 0.1 and 0.2, respectively. SNP counts are shown in the legend after the dash. B. Density profiles show variants of ClinVar, somatic mutations (COSMIC) and 1KGP database in coding regions. C. Densities of damaging and benign variants predicted by Polyphen-2 based on the HumanVar database and annotated on 1KGP database by ANNOVAR software.

We also compared Pref and Palt for different types of single nucleotide variants (SNVs) in coding (Figure 7B) and non-coding regions (Supplementary Figure S5). Clinically relevant mutations from the Clinvar database are almost indistinguishable from 1KGP in coding regions and indeed a Kolmogorov–Smirnov (KS) test gives a p-value of 0.18 showing an insignificant difference (see Supplementary Table S6). On the contrary, somatic mutations have a clear tendency to mutate towards a more probable base (Palt > Pref) supported by a p< 10^−15^ in the KS test. In non-coding regions, the somatic mutations are also shifted towards a higher probability for the alternative and have the same peak at high reference probability as 1KGP.

To see if there is a difference between damaging and benign SNPs, we show the same densities for Polyphen2 predictions [17] on Chr1 in Figure 7C. On Chr1 there is a total of 32,841 SNPs classified as benign and 15,299 SNPs classified as damaging. There is a small, but significant (KS test (p< 10^−15^, see Supplementary Table S6)), shift of the damaging SNPs towards higher probability of the alternative allele. We saw that for only 21% of damaging SNPs the reference allele had the highest probability whereas for 29% the alternative allele had the highest probability. For benign SNPs, these numbers are 26.5% and 24%. This difference is highly significant (Chi-squared test p≃ 10^−9^, see Supplementary Table S7).

### Context-dependent models of substitutions

It is possible to estimate context dependent models of single nucleotide substitutions from a set of known variants. Since SNV sampling is very biased and variants are not fully observed, the context size needs to be much smaller than for the nucleotide distribution models described above. In the previously mentioned work [5] a seven nucleotide context is used. Here we want to explore the possibility of using our genome models to obtain models of substitutions. The rationale is that to maintain the context dependent nucleotide probabilities, they must be reflected in the mutability.

We assume the genome has reached approximate equilibrium. To keep this state, the mutability towards a nucleotide should be higher, the higher the probability of that nucleotide is in the given context. Therefore we set the probability of a mutation from *a* to *b* to be proportional to the probability of nucleotide *b* (in that context) with a constant that depends on the nucleotides and which can also depend on the context. This model is inspired by the general time-reversible stationary Markov model [18, 19], in which the off-diagonal rates are *µ*_*ab*_ = *α*_*ab*_*π*_*b*_ with symmetric *α*_*ab*_ for nucleotides *a* ≠ *b* and the equilibrium distribution *P*(*a*) = *π*_*a*_. The mathematical theory does not apply directly here, because reversibility is too restrictive, so we do not require the *α* matrix to be symmetric, but we can still estimate an *α* matrix that best fits a set of variants. For lack of a better term, we call *α* the “alpha matrix”.

Whereas the nucleotide distribution can be estimated from the whole genome using large contexts, the *α*s must be estimated from observed mutations. We hypothesize that the *α*s are less context dependent, and thus can be estimated from a smaller context than the nucleotide distributions. Details of the estimation procedure is described in Methods.

We estimated *α*s from all chromosomes except Chr1 for symmetrical contexts of size 0, 3, 5, and 7 (*k* = 0, 1, 2, and 3) using SNPs from the 1KGP and the BM14 model for the nucleotide distribution. The alpha matrix is shown in Table 1 (left) for *k* = 0. Notice that it is essentially strand-symmetric, but not symmetric in normal matrix-sense, so it violates reversibility. Similarly, we estimated a simple conditional model with a 7-mer context (*k* = 3) from the same data, which is called the simple model in the following. The simple model is similar to one of the models in [5], but the variants used for estimation are slightly different. The models were then applied to Chr1 where we calculated the probability of a mutation given the context for all positions with an observed SNP. The total fraction of sites with probability above 0.25 is very small for all models, see Figure 8A. In Figure 8B the fraction of sites with a certain mutability that has an observed SNP is plotted against mutability for some of the models. Ideally these should be linear, but we see a significant deviation from linear for the simple model and for the *α* models with *k* > 0. The models with *k* = 1–3 behave almost the same, and up to a substitution probability of ∼ 0.25 they are very close to the simple model.

**Table 1.**
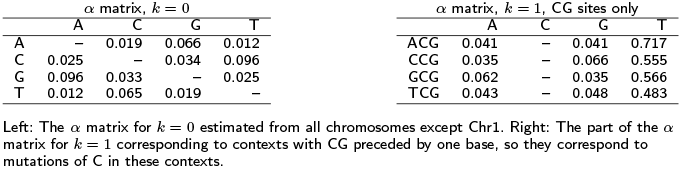
*α* matrixes for *k* = 0 and *k* = 1 estimated by substitution model.

| $\alpha$ matrix, $k=0$ | | | | | $\alpha$ matrix, $k=1$, CG sites only | | | | |
| --- | --- | --- | --- | --- | --- | --- | --- | --- | --- |
|  | A | C | G | T |  | A | C | G | T |
| A | — | 0.019 | 0.066 | 0.012 | ACG | 0.041 | — | 0.041 | 0.717 |
| C | 0.025 | — | 0.034 | 0.096 | CCG | 0.035 | — | 0.066 | 0.555 |
| G | 0.096 | 0.033 | — | 0.025 | GCG | 0.062 | — | 0.035 | 0.566 |
| T | 0.012 | 0.065 | 0.019 | — | TCG | 0.043 | — | 0.048 | 0.483 |

Left: The $\alpha$ matrix for $k=0$ estimated from all chromosomes except Chr1. Right: The part of the $\alpha$ matrix for $k=1$ corresponding to contexts with CG preceded by one base, so they correspond to mutations of C in these contexts.

**Figure 8.**
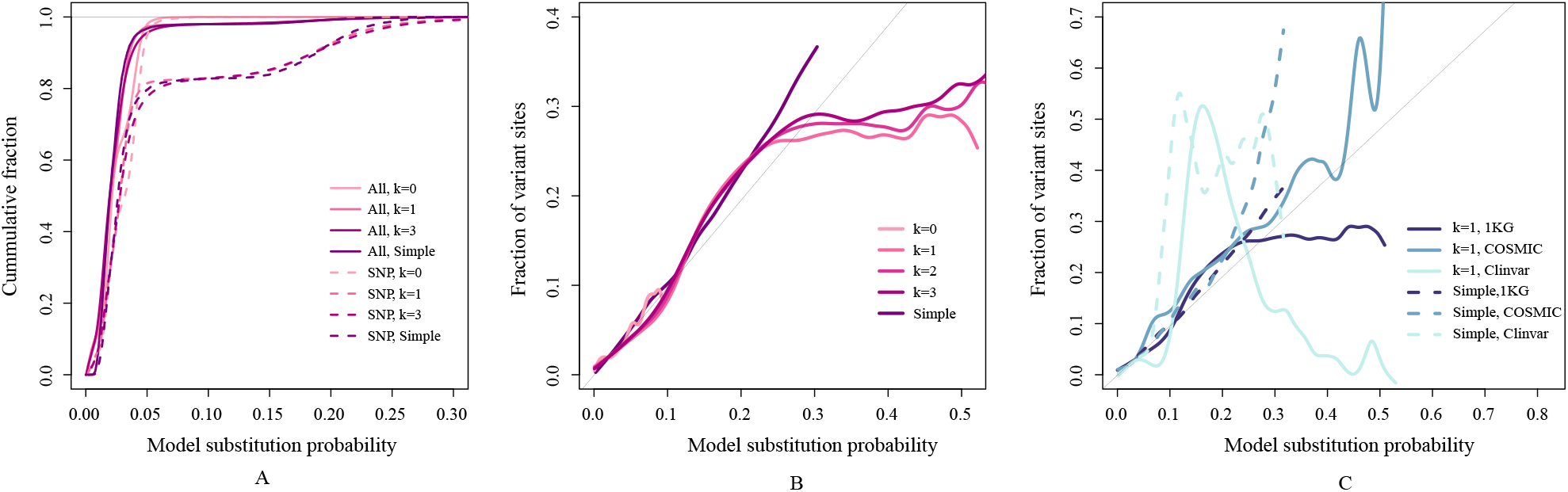
Substitution model. Model substitution probabilities shown for the models with context-insensitive *α* (k=0), the ones with *α* depending on 1, 2, and 3 bases to each side (k=1, 2, 3), and the simple model conditioned on the 3 bases to each side. The model substitution probability for a site is the sum of the probabilities for the three possible substitutions. A. The cumulative distribution of model substitution probabilities for all sites (solid lines) and for SNPs (dashed) on Chr1 shown for the five models. Note that for all models there are very few sites with substitution probability above 0.3. B. The fraction of sites on Chr1 with an observed variant in the 1000 Genomes project (1KGP) plotted against p. The y values are SNP counts in small probability intervals (10^−4^) divided by total counts. The curves are smoothed with splines. Estimates are noisy for larger probabilities due to low counts. C. As B for SNPs in 1KGP, Clinvar and COSMIC for the *k* = 1 model and simple only. For latter two, counts are scaled so they sum to the number of SNPs in the 1KGP set for Chr1. For high mutability values there are few SNPs, so the curves are very noisy especially for Clinvar.

Above a mutability of 0.25, our models with *k* > 0 deviate significantly from the diagonal line. It turns out that these rare reference genome sites with high substitution probability are mainly CpG sites. The alpha matrix for *k* = 1 is shown in Table 1 for the CG contexts, where it is evident that the C to T values are very large, ranging from 0.48 to 0.72, which should be compared to the largest *α* of 0.22 that is not a CG context, see (Supplementary Table S8). For contexts where the T has high probability according to the nucleotide distribution, the substitution probabilities will become large, because it is the product of *α* and the nucleotide probability. It suggests – as expected – that these substitutions are very likely at unselected positions.

We applied the model also to SNVs from Clinvar and COSMIC as shown in Figure 8C for *k* = 1 and for the simple model. The number of variants with mutability values above 0.3 for the *k* = 1 model is relatively small. For Clinvar only 296 SNVs out of 42000 have a mutability larger than 0.3 and for COSMIC this number is 2760 out of 120000. It means that the data are noisy as seen in Figure 8C, but it is evident that the somatic SNVs from COSMIC follow the model more closely than germline SNPs in this domain.

## Discussion

We developed context dependent models of the nucleotide distribution in the human genome. The most advanced one, a bi-directional Markov model with a context of 14 nucleotides to each side, can predict a nucleotide with 51% accuracy. We use interpolation from lower orders, so it is in principle possible to go above *k* = 14, but we saw that this did not change the model very much, and the predictability of just above 50% is close to an upper limit for this type of model.

In this work our objective has been to apply simple interpretable models to the problem. Previous studies have applied neural networks to the human genome by sequence context to obtain DNA representations for other tasks. This has been used for prediction of the effect of non-coding variants [20] and the regulatory code of the accessible genome [21], for instance. The DNAbert model [22] is more related to the present work. It is a transformer neural network, which in the pretraining is trained to predict k-mers (k=3-6) from the surrounding sequence context. However, the focus is on using it for other prediction tasks, and direct comparison to our models is not possible. We have used neural networks ourselves for the same task for prediction of bases from the context [23]. Using a larger context in the neural network leads to marginally better prediction accuracy, but more importantly differences in performance depending on context.

The high predictability of our model is, to a large extent, due to repeats. It is interesting that approximately half the human genome is said to be repetitive [24], which superficially coincides with the predictability, but an exact definition of repetitive regions is a challenge and some report a higher repetitive fraction (see e.g. [25]). For *A. thaliana* and *C. elegans* the predicability was 40% and 45%, respectively, and they both have 12-13% repeats [26], and although the model was of lower order, it suggests that predictability could be used as a measure of the repetitiveness of a genome. This, however, would require more extensive analyses.

Not surprisingly, the predictability is highly dependent on the type of the genomic region. Coding regions can be predicted with only 36% accuracy, whereas Alu repeat regions are at 87% and simple repeats even higher (Figure 3). When looking more closely at splice sites we see – as expected – a negative correlation between conservation and the probability of the reference base (Figure 5), although such a correlation is weak, when looked at genome wide due to the lack of conservation of repeats. There are also differences between chromosomes, where especially the Y chromosome and Chr19 stand out with higher predictability than others, which is likely due to their high repeat content.

The model was applied to the genomes of *Arabidopsis thaliana, Caenorhabditis elegans, Escherichia coli*, and *Saccharomyces cerevisiae*. Due to the smaller genome sizes a bidirectional Markov model with *k* = 10 was used. The large differences between species observed is an indication of quite different composition of genomes. Interestingly some species have two peaks in the density of the reference probability, which is partly explained by differences between coding regions and non-coding.

We compared the probability of the reference allele to the alternative allele on single nucleotide variants from the 1000 Genomes Project. There is a peak with SNPs that have a reference probability close to one, which skews the distribution away from symmetry (Figure 7A). Almost all SNPs in this peak (with reference probabilities over 0.875) fall in repeat regions and one possibility is that some of them are mapping artefacts. They also have relatively low allele frequencies, and when considering only SNPs with high allele frequency, the plot becomes symmetric. Therefore, another factor that may explain the asymmetry is that the reference genome, which is not a genome of a single individual, contains very few rare alleles.

The difference between the probability of the reference allele and the alternative allele for coding SNVs in the 1000 Genomes Project was compared to SNVs from somatic mutations and clinically relevant SNPs from Clinvar (Figure 7B). Here we see a statistically significant shift of somatic SNVs towards higher probability for the alternative allele, which suggest that somatic mutations tend to favor more probable bases. Similarly, we see a significant difference between damaging and benign SNPs (as classified by ANNOVAR) as seen in Figure 7C. Surprisingly, the damaging SNPs seem to have a higher probability according to our model than benign ones.

The sequence models presented here estimate distributions of the bases for a given context and reflect inherent properties of the cellular machinery responsible for replication, error correction, and so on, as well as the physical properties of DNA, such as curvature and bendability. A mutation that moves a base closer to this distribution is likely to be more probable than one that moves it away, at least if selection is ignored. To explore this, we have derived a model that takes the context dependent nucleotide distribution into account.

In our model, we are assuming that the variation of a site in the human DNA can be described by a context sensitive continuous Markov model with a rate matrix that is a product between the nucleotide distribution and an “alpha matrix”. The alpha matrix can be estimated from known variants and it can depend on a smaller context than the model for the nucleotide distribution and can be estimated from a relatively small number of SNVs. It means that our model for mutability have a very large context due to the context dependent nucleotide distribution even if the alpha matrix uses a smaller context.

The model does not depend strongly on the context size for the alpha matrix for contexts of the two neighbours or larger (*k* ≥ 1). Our models behave very similarly to a simple mutability model, which is estimated from SNPs alone and a context of three nucleotides to each side except in a regime of very high mutability (Figure 8B). Our models seem to over-estimate the SNP mutability from 1KGP when the values are larger than about 0.25. However, this is not the case for somatic mutations, and the mutations seem to be well-described by these models (Figure 8C).

The model is inspired by the general time-reversible model from evolutionary theory, which has six free parameters corresponding to a symmetric alpha matrix, and with rates depending on the equilibrium distribution. However, although time-reversibility would be desirable, it is not likely that the context dependent nucleotide distribution we estimate is an equilibrium distribution for the entire genome. In fact, when inspecting the estimated alpha matrix for zero context (Table 1) and a context of one nucleotide to each side (Supplementary Table S8), it is evident that it is not symmetric. For the latter there are very large deviations from symmetry for contexts with NCG, where N can be any base. In these contexts, *α*_*CT*_ is consistently 10-20 times larger than *α*_*T C*_ corresponding to a strong tendency to mutate from CG to TG.

Even if the *α* matrix depends on a small context, the substitution still depends on the full context of the nucleotide distribution. This construction is very attractive, because substitution models estimated from variants alone need to have small contexts due to the limited number of variants and the strong sampling biases.

## Conclusions

There are strong context dependencies of nucleotides in genomes. We have shown how one can estimate a model of the nucleotide probabilities depending on contexts up to 14 nucleotides to each side. Building on these models, it was shown how it is possible to make models of mutations that combine the context dependent nucleotide probabilities with a mutation matrix, called the alpha matrix, to give mutation probabilities (“mutabilities”) that depend on the same large context. It was shown that these models fit observed mutations very well and especially somatic ones. Importantly, the alpha matrix can depend on a much smaller context of just one to three bases to each side and does not depend strongly on this parameter. These models can form the basis for a better understanding of human mutations and we believe it will be possible to use them in a wide range of applications from GWAS studies to analysis of somatic mutations.

## Methods

### Conditional probability models for the central base

The base at position *i* (chromosome, coordinate) in the reference genome is called *x*_*i*_ and the symmetric sequence context around it is called

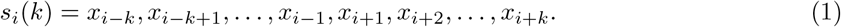

If it is clear from the context which *k*, we call it *s*_*i*_ to ease notation. To estimate the conditional probability of base *b* at position *i*, we use the counts *n*(*b*|*s*_*i*_) of the occurrences in the same context throughout the reference genome (on both strands):

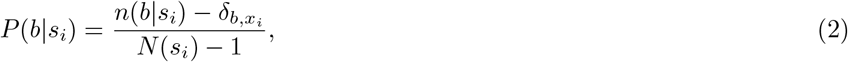

where

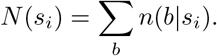

We use the Kronecker 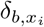, which is 1 if *x*_*i*_ = *b* and otherwise 0, to ensure that we only count *other* contexts, when estimating probabilities at position *i*. This is leave-one-out cross-validation and is discussed further below.

For large contexts, the counts become small and thus the probabilities cannot be reliably estimated. To interpolate between different orders of the model, we use regularization by pseudo-counts obtained from the *k* − 1 model. Specifically, for order *k*, we define pseudo-counts

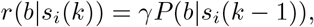

where *γ* is the strength of pseudo-counts. Now the model of order *k* is estimated as before, but using the actual counts plus pseudo-counts,

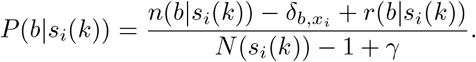

The advantage of pseudo-counts is that they have minor influence, when there is plenty of data (actual counts are high), but have strong effect at low counts. With *k* = 4 counts are on average 6 *10^9^*/*4^9^ ≃ 23000, so we assume that psudo-counts are not needed. Therefore, our interpolated model starts with unregularized estimates for *k* = 4, and then use the pseudo-counts iteratively for *k* = 5 to *k* = 7 for the interpolated model. We used a strength of *γ* = 100 for the pseudo-counts (a few experiments showed that the model is relatively robust to changes in *γ*, see below).

### Markov models

In a Markov model of order *k*, the probability of a base is conditioned on the *k* previous bases. If we redefine the *k*-context in (1) to be the *k* previous bases,

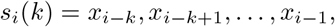

we can use exactly the same formulation as above. In this case however, the context size is not 2*k* letters as above, but only *k* letters. Therefore, one can estimate Markov models up to sizes around *k* = 14 for the human genome, and we used a model interpolated from *k* = 8 to *k* = 14 analogously to the central interpolated model described above.

Due to the interpolation, larger *k* are possible, and we performed a small experiment with *k* ranging from 10 to 20 and with four different values of the interpolation constant *γ* resulting in Supplementary Figure S2. These tests were done only on chromosome 20 with a model estimated from all chromosomes *except* 20. Although small gains can be obtained with larger *k* values and different *γ*, we decided to stick to our initial choice of *k* = 14 and *γ* = 100.

Estimating a “forward” Markov model from both strands of the human genome will automatically make it strand-symmetric. For a given position in the genome, the model can therefore give two sets of base probabilities: one for the forward strand and one for the reverse strand. Our final Markov probabilities are the average between the two as described in the main text and referred to as bidirectional.

### Cross-validation

Our way of estimating the conditional probability of seeing one of the four bases given the surrounding context can be seen as a leave-one-out procedure. In particular, the estimate depends on the reference base at the considered position as well as the context. To obtain an estimate that is independent of the reference base at the position, a natural way to proceed is to consider the average of the four base-dependent estimates over all occurrences of the given context. This average turns out to be equal to the estimate that includes all positions. To see this, average (2) over all sites (skipping the *k* dependence for clarity) gives the probability of a base *b*:

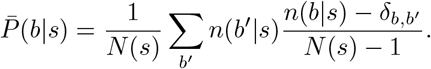

Here the base we are summing over is called *b*′ to distinguish it from the base *b* in question. Since Σ_*s*_*n*(*b*′|*s*)*δ*_*b,b*′_ *= n*(*b*|*s*), we get

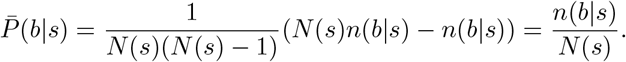

We also assessed our models by cross-validation by chromosomes. One chromosome was used as test data, and the remaining chromosomes as training data. We repeated this step 24 times to calculate the fraction correct predictions for each chromosome.

### Substitution models

A simple model estimates mutability as the fraction of all sites with context *ŝ* having a specific mutation. More specifically,

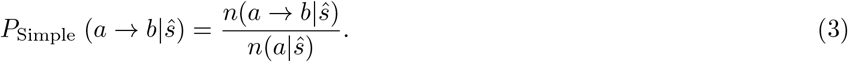

Here *n*(*a* → *b*| *ŝ*) is the number of observed mutations *a* → *b* in context *ŝ* and *n*(*a*|*ŝ*) is the number of times we see reference base *a* in context *ŝ* (as above). We use *ŝ* to indicate that the context may be different from the context *s* for the genome model above. We have used this model with a symmetric context of three bases to each side, which we call the simple model.

We will now derive a continuous time Markov model with context dependent substitution rates *µ*_*ab*|*s*_ that takes the nucleotide distribution into account. We also assume a constant evolutionary time, which is infinitesimally small compared to the rates, so we can approximate the substitution probability by the first-order term in the Taylor expansion of an exponential

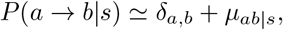

where time is set to 1. The diagonal rates are −Σ_*b*≠*a*_ *µ*_*ab*|*s*_, so in the following we will not write the diagonal terms. For a stationary, reversible Markov model with *P* (*a*|*s*) as equilibrium probabilities the rates can be written as

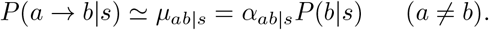

with a symmetric matrix *α*_*ab*_. This is the general time-reversible six-parameter model (see e.g. [19]). Inspired by this model, we assume that mutability is given by the same equation, but without requiring that the nucleotide distribution is the equilibrium distribution and without requiring that *α* is symmetric.

The above expression factorizes the rates into the nucleotide distribution and the *α*-term that encapsulates the mutations. Now we assume the *α*s depend on a *smaller* context *ŝ* than the context *s* for the genome model *P* (*a*|*s*), so the above can be written as

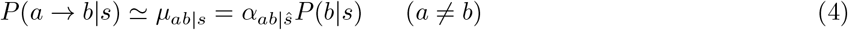

In analogy with (3), *P* (*a* → *b*|*s*) = *n*(*a* → *b*|*s*)*/n*(*a*|*s*) with *s* instead of *ŝ*, so combining with the above

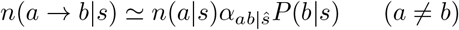

To estimate the *α*s we sum over all contexts that contains *ŝ*, which we write as *s*| *ŝ* ⊆ *s*, so

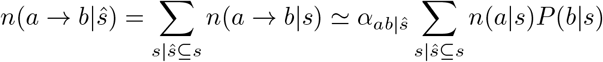

The last sum depends only on the nucleotide distribution. It can be rewritten as a sum over all positions in the genome, where the reference base, *r*_*i*_, equals *a* and where the context is *ŝ*. We call this term *Z*_*ab*| *ŝ*_,

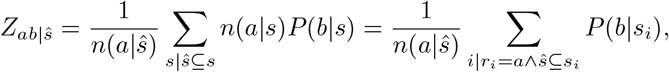

For convenience, it is normalized by *n*(*a*| *ŝ*), so it is the average probability of base *b* over all positions with reference base *a* and context *ŝ*. As an estimate of *α* we then have

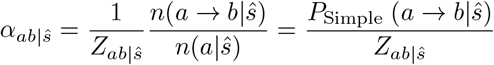

Note that we can rewrite the the original probability (4) in terms of the simple model as

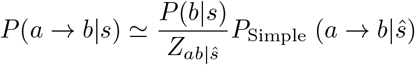

for *ŝ* ⊆ *s*. The factor is 1 when *ŝ* = *s*, so the models are identical as they should be when they use the same context. The equation directly shows how the wider context from the genome model can modulate the simpler estimate. If the probability of base *b* in context *s* is larger than the mean *Z*_*ab*|*ŝ*_, the mutability becomes larger than in the simple model, and if it is smaller, the mutability becomes smaller.

The first order approximation assumes the rates are small. When calculating the total mutability of a site, we therefore use the approximation 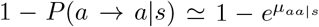. For small *α*’s it makes little difference whether it is the exponentiated form or not.

### Data

The human reference genome, GRCh38.p13, was downloaded from NCBI (released March 2019 by Genome Reference Consortium). We considered only primary assemblies of chromosomes 1 to 22 and X, Y. Genomic annotation bed files were downloaded from UCSC Table Browser. These are 3’-UTR, 5’-UTR, CDS, Introns, Genes, and Repeats. Conservation scores file (PhastCons100way) was downloaded from the UCSC as well.

Variants were downloaded from the 1000 Genomes project (released March 2019, phased 20190312_biallelic_SNV_and_INDEL) in VCF format. The INDELs were filtered from 1KGP dataset.

ClinVar (clinvar_20200310.vcf) [27, 28] and somatic mutations (CosmicCodingMuts.vcf and CosmicNonCodingVariants.vcf) [29] data were obtained from NCBI and COSMIC, respectively.

The genomes and GFF files of *Arabidopsis thaliana* (TAIR10.1), *Caenorhabditis elegans* (WBcel235), *Escherichia coli* (str. K-12 substr. MG1655), *Saccharomyces cerevisiae* (R64) were downloaded from NCBI.

### Data analysis

#### Model implementation

Counting of k-mers and estimation of probabilities is implemented in the C programming language. The program counts the contexts for each site using a Burrows-Wheeler transform (BWT) [30] rather than storing the k-mers, because it is much more efficient for the interpolated models. The program is called predictDNA and relies on an index built with the program makeabwt.

One program, called makeabwt, is used for construction of an index from a fasta file containing the genome sequences. If there are multiple sequences, they are concatenated with termination symbols in between and the suffixes are sorted. The BWT is constructed from the sorted suffixes and saved. An FM index [31] is constructed to ease the search of the BWT. To limit memory usage, the values are stored in first-level checkpoints for every 2^1^6 positions as long integers (8 byte) and for every 256 positions the difference from the nearest first-level checkpoint is stored as a short integer (two bytes). We used an index containing both the forward and reverse complements strands of the genome.

Another program, called predictDNA, use the index to look up *k*-mers. This is done using the standard backward search of the BWT/FM-index [31]. The size of the resulting suffix interval equals the number of the *k*-mers in the genome and these are used for calculating the conditional probabilities.

The advantage of using a BWT is that the index can be used with any *k* and thus facilitates the interpolated models. An naive approach using table-lookup would require a new table for each value of *k* and a table of 4^15^ ≃ 10^9^ integers for *k* = 14, which corresponds to 4GB of memory and this would become 16GB for *k* = 15, etc. The index used for this work use around 8GB of memory.

#### Model Performance

We calculated the probabilities of the four bases for every position in the human genome using the software predictDNA we developed. We tested different *k*’s, but used the same interpolation constant, *γ* = 100, for all models. We counted the correct sites for which the reference alleles gave the highest probabilities of the four bases, to calculate the fraction correct for each chromosome.

Furthermore, we overlapped the bed files with models’ outputs via bedtools [32, 33] to get the feature-specific fraction correct and predicted probabilities. These were used to obtain the performance of our models for different regions of human genome.

Based on CDS bed file and human genome fasta file, we calculated average probabilities for the positions around the human 3’ and 5’ splice sites. We included 500 nucleotides beforer and 100 after the 3’ splice site and, similarly, 500 before and 100 after the 5’ splice. Besides, we extracted the conservation scores of Phast-Cons100Way for the same regions [34]. Those results were shown in Figure 5.

#### SNP Variants Analysis

We kept only single nucleotide bi-allelic variants in 1KGP, ClinVar and COSMIC databases for the following analysis, and we filtered INDELs. Based on central model and BM14 results, reference and alternative allele probabilities for each SNP sties in these three databases were extracted. The triangle plots (Figure 6) were made by using reference probabilities against alternative probabilities of all SNPs in 1KGP database.

In order to understand the possible asymmetry shown by the cluster of many sites in the corners of the triangle plot, we separated SNPs with allele frequency greater than 0, 0.01, 0.1 and 0.2. To present the different types of SNPs in coding and non-coding parts, we did the density plots also by using Pref minus Palt for SNPs in 1KGP, ClinVar and COSMIC databases. Additionally, we used ANNOVAR software[35] to annotate benign and damaging SNPs on 1KGP, which were predicted by PolyPhen2 [17]. These are sites associated with single genetic disease.

We developed the subsitution model to estimate the mutability of SNVs as described above. We estimated the *α* matrix for *k* = 0, 1, 2, 3 for all SNPs 1KGP outside of Chr1. The model was applied to chromosome 1, where we calculated the probability of a mutation from the BM14 and the alpha matrices. These were compared to observed SNVs in 1KP, ClinVar, and COSMIC on Chr1.

#### Test Bi-directional Markov Model on Other Species

The bi-directional Markov model with was tested on the chosen species and also human genome. We used *k* = 10, *γ* = 100, and interpolated from *k* = 6, instead of using the same parameters as BM14, that is because of the smaller genome size of these species. The densities of the reference base probabilities were plotted (Supplementary Figure S4A). We separated the CDS and non-coding regions of A.*thaliana*, C. *elegans* and S. *cerevisiae* according to the GFF files and made a density plot to show the distributions of CDS and non-coding of these three species.

### Software

Our software is open source and available at GiHub: https://github.com/AndersKrogh/abwt/releases/tag/v1.2.1a. We wrote several scripts in Perl and Python for data analysis and these are all available in the GitHub release. The usage of these scripts is described in README files. All the figures made in R and this code is also available.

## Abbreviations

BM14: Bidirectional Markov model with 14 bases as context
p.p.: percentage points
CDS: Coding Sequence
Chr: Chromosome
Pref: Probability of reference
Palt: Probability of alternative
1KGP: 1000 Genomes Project
SNP: Sigle nucleotide polymorphism
SNV: Single nucleotide variants
BWT: Burrows-Wheeler transform

## Declarations

## Acknowledgements

We thank Hanne Munkholm for her big help and support with compute servers.

## Availability of data and materials

All data used in this study are publicly available. All data can be downloaded from NCBI, UCSC, 1KGP and COSMIC database as we mentioned in our methods.

The links to the genomes of the species we used:

*Homosapiens* (https://www.ncbi.nlm.nih.gov/genome/?term=GRCh38.p13), *Arabidopsisthaliana* (https://www.ncbi.nlm.nih.gov/genome/?term=TAIR10.1), *Caenorhabditiselegans* (https://www.ncbi.nlm.nih.gov/genome/?term=WBcel235), *Escherichiacoli* (https://www.ncbi.nlm.nih.gov/genome/?term=Escherichia+coli),

*Saccharomycescerevisiae*(https://www.ncbi.nlm.nih.gov/genome/?term=Saccharomyces+cerevisiae) The CDS, Introns, 3’-UTR, 5’-UTR, Genes, Repeats and Conservation score are download from UCSC Table Browser (https://genome.ucsc.edu/cgi-bin/hgTables)

1000 Genomes Project (http://ftp.1000genomes.ebi.ac.uk/vol1/ftp/data_collections/1000_genomes_project/release/20190312_biallelic_SNV_and_INDEL/)

clinvar_20200310 was used for Clinical SNPs analysis (https://ftp.ncbi.nlm.nih.gov/pub/clinvar/vcf_GRCh38/archive_2.0/2020/)

Coding and non-coding mutations of COSMIC (https://cancer.sanger.ac.uk/cosmic/download)

## Ethics approval and consent to participate

Not applicable.

## Consent for publication

Not applicable.

## Authors’ contributions

AK and PF initiated the project. YL and AK performed most analyses and drafted the paper with assistance from CG and PF. All authors participated in revision and approved the final version.

## Additional Files

Additional file 1 — Supplementary tables S1-S8

Supplementary tables: Table S1, S2, S3, S4, S5, S6, S7, S8

Additional file 2 — Supplementary Figures

Supplementary figures: Figure S1, S2, S3, S4, S5

