## Supplementary Figures for "Context dependency of nucleotide probabilities and variants in human DNA"

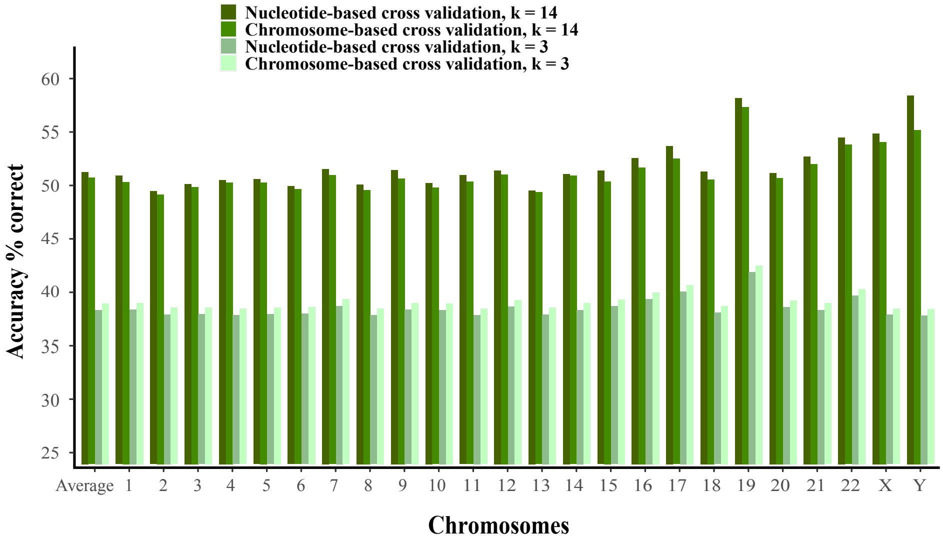


Figure S1: Chromosome based cross validations for baseline model and Bidir-Markov model. For each chromosome, the overall prediction accuracy is calculated for a model is estimated from the other chromosomes (chromosome-based cross validation). The overall average is weighted by chromosome sizes. These are compared to the nucleotide-based cross validation accuracies used in Figure 1.

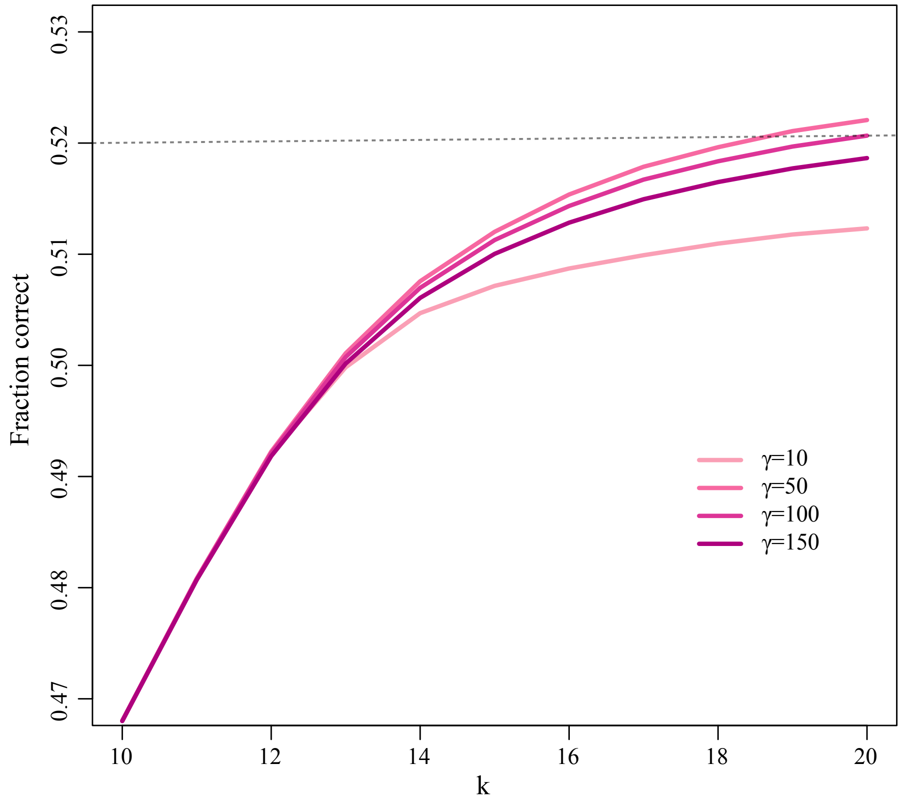


Figure S2: Accuracy of the bi-directional $k$-th order Markov model for different strengths of regularization, $\gamma$. Results are shown only for Chromosome 20 with the model estimated from all the other chromosomes.

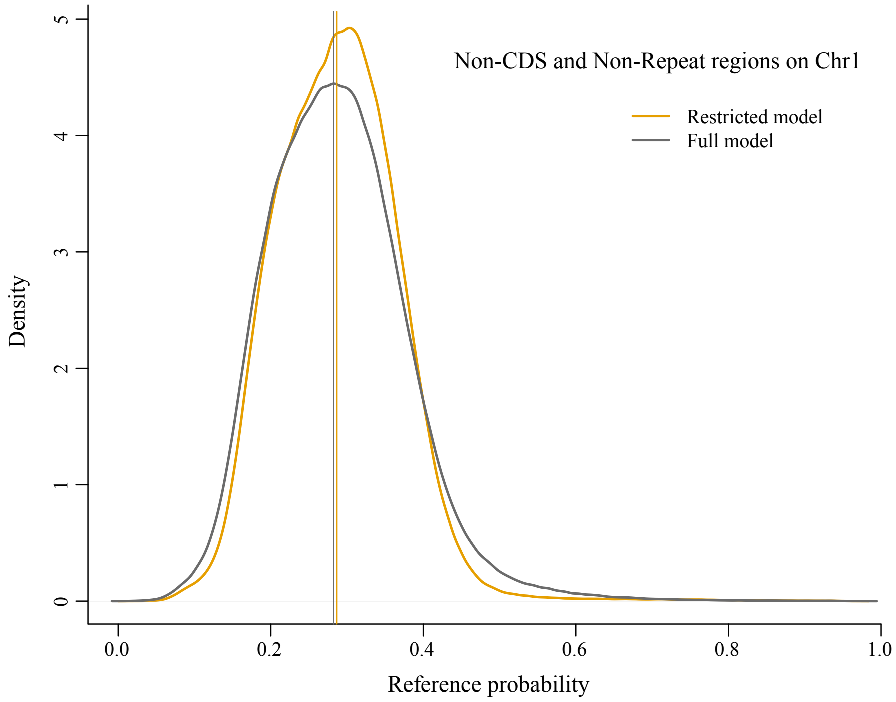


Figure S3: Comparison of restricted and full model based on density profile of reference probabilities. Density profile of the reference probabilities for the full model was shown as a dark grey line and the other for a model estimated on non-repeat and non-coding regions on Chromosome 1. The yellow and gray vertical lines represent the median probabilities of restricted model and full model, which are 0.286578 and 0.282368, respectively.

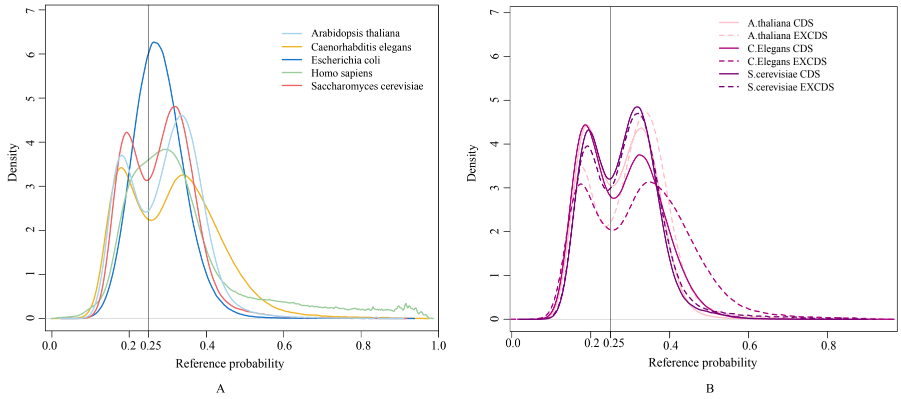


Figure S4: Density profile of reference probabilities of different species. A. Those species were estimated via 10-$k$ context bidirectional Markov model, $\gamma$ = 100 interpolated from 6. B. Density plots of CDS regions and non-CDS for the species, which have two peaks in Figure S4A.

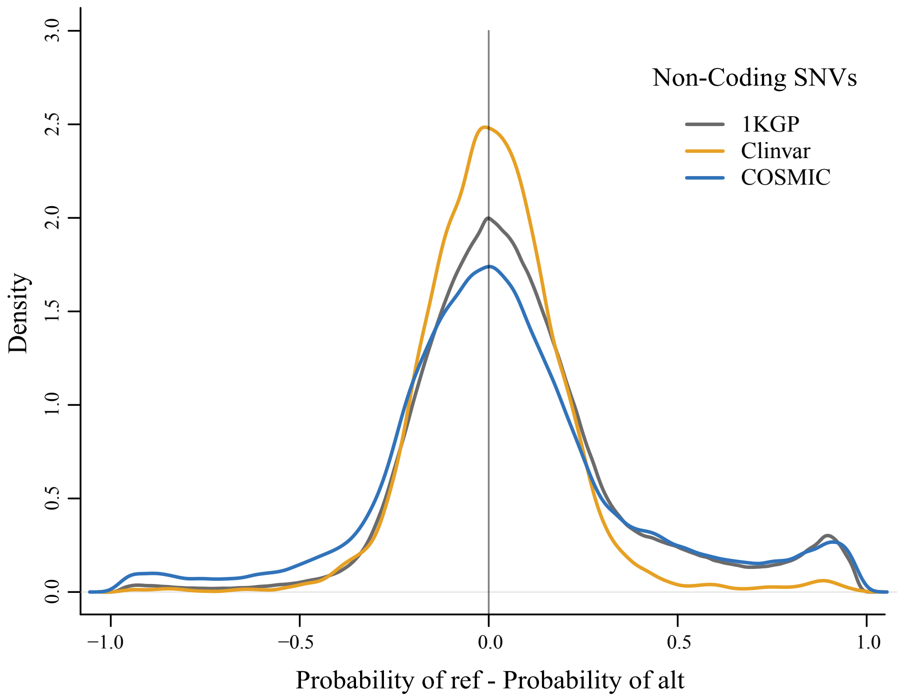


Figure S5: Density profiles of Pref - Palt for SNPs on Chromosome 1. Density profiles show ClinVar, somatic mutations (COSMIC) and 1KGP SNPs in Non-Coding regions, respectively.
